# Determining a reasonable range of relative numerical tolerance values for simulating deterministic models of biochemical reactions

**DOI:** 10.1101/256214

**Authors:** Ming Yang, Louis Z. Yang

## Abstract

What values of relative numerical tolerance should be chosen in simulation of a deterministic model of a biochemical reaction is unclear, which impairs the modeling effort since the simulation outcomes of a model may depend on the relative numerical tolerance values. In an attempt to provide a guideline to selecting appropriate numerical tolerance values in simulation of *in vivo* biochemical reactions, reasonable numerical tolerance values were estimated based on the uncertainty principle and assumptions of related cellular parameters. The calculations indicate that relative numerical tolerance values can be reasonably set at or around 10^−4^ for the concentrations expressed in ng/L. This work also suggests that further reducing relative numerical values may result in erroneous simulation results.

---

A deterministic model of a biological process in the form of differential equations presents mathematical relationships among variables of precise values, but in simulation of such a model, only approximate values of the variables can be employed. The approximation is governed by either an absolute or a relative numerical tolerance value. An absolute numerical tolerance value is the absolute difference between the actual and expected values. Dividing such an absolute difference by the expected value gives rise to a relative numerical tolerance value. Because a model of a biological system may produce different simulation outcomes with different ranges of numerical tolerance values, it is important to determine a reasonable range of numerical tolerance values. Currently a reasonable range of numerical tolerance values for modeling a biological process such as that involving biochemical reactions has not been explored

Numerical tolerance issues have been studied before, which led to the general conclusion that decreasing numerical tolerance values to a sufficiently small and yet practical level may effectively correct erroneous outcomes in computer simulation (Byrne and Thompson, 2013). The erroneous outcomes arise from numerical stiffness, a phenomenon of unacceptably large errors in dependent variables as a result of small errors in independent variables (DiStefano III, 2015). Here we present a case that a stable numerical outcome may be achieved only when the numerical tolerance values approach zero. This case was previously described in an attempt to model oscillations of biochemical reactions in a negative feedback system (Yang and Yang, 2015). In this case, the magnitudes of the oscillations of chemicals *x* and *y* decrease along with the reduction in numerical tolerance values, but chemical concentrations remain oscillatory even when the relative and absolute numerical tolerance values are reduced to 10^−10^ and 10^−12^, respectively (Fig. 1). It seems that the chemical concentrations will remain oscillatory unless the numerical tolerance values approach zero or are much smaller than those in Fig. 1C. A question then arises: which simulation outcome should be accepted in this case, an outcome with a certain range of numerical tolerance values far from zero or one with the numerical values approaching or close to zero?

**Fig. 1.**
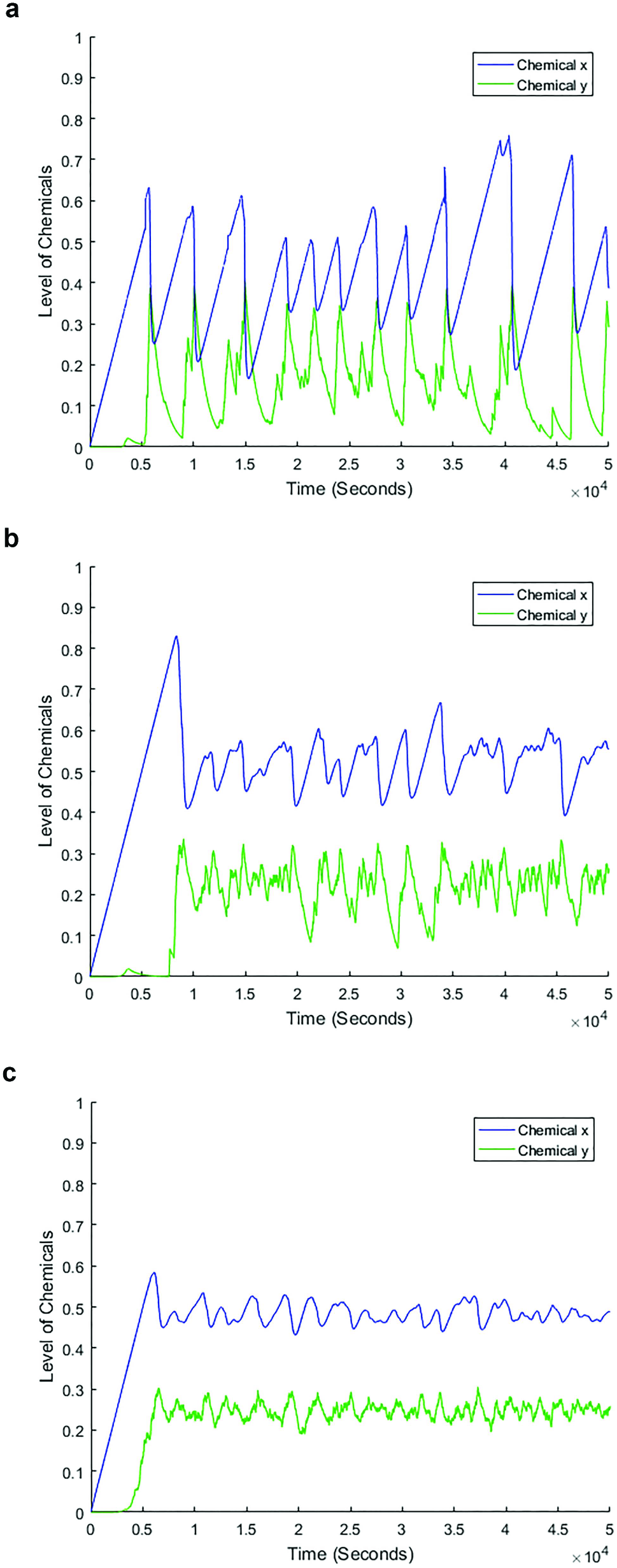
Simulation outcomes of a negative feedback system of biochemical reactions at three numerical tolerance levels. This is a two-ODE system numerically integrated as described for Fig. 2B in Yang and Yang (2015). (a) Relative numerical tolerance = 10^−3^, and absolute numerical tolerance = 10^−6^. (b) Relative numerical tolerance = 10^−5^, and absolute numerical tolerance = 10^−8^. (c) Relative numerical tolerance = 10^−10^, and absolute numerical tolerance = 10^−12^.

We would argue that the answer to the above question is an outcome with the numerical tolerance values far from zero. We think that at the fundamental level, chemical concentrations cannot be precisely determined and the inherent variance of a chemical concentration is governed by the uncertainly principle. The uncertainty principle was typically applied to a subatomic particle. Because each molecule can be considered as a population of subatomic particles, the uncertainty level of such a population in position and energy state should be equal to or more than that of a subatomic particle, depending on the complexity of the molecule. The uncertainty level of a subatomic particle, therefore, can serve as a guidance value for the estimation of the lower limit of the uncertainty level of molecules. Precisely calculating the uncertainty level of a molecule, or a population of molecules in the case of determining the concentration of a molecular species, is challenging due to the complexity arising from the large number of subatomic particles involved and the dynamics of the molecule dictated by its internal structure and external environment. Towards finding the lower limit of the uncertainty level of a population of molecules, we start from the uncertainty level of a subatomic particle, which is typically presented as the following:

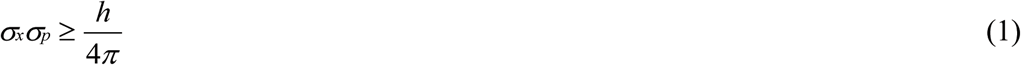

where *σx* and *σp* are the standard deviations of the position and momentum of a particle, respectively, and *h* is the Planck constant. Now let’s consider a biochemical reaction as a representative case. The kinetics of a chemical reaction is typically described by an equation with variables of the concentrations of the reactant(s) and product(s), and the standard deviation of the concentration *σc* is, in effect, the relative numerical tolerance value, which exerts its influence on the simulation when the concentration values are reasonably large (e.g. >0.1). The standard deviation of the concentration of a chemical, *σc*, can be calculated as the following:

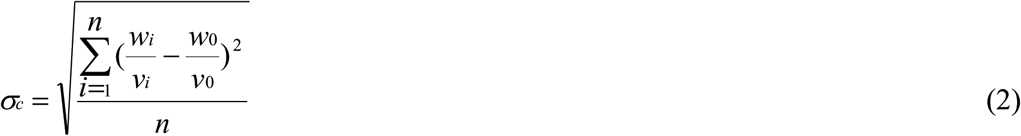

where *w*0, *v*0, and *n* are the mean of the chemical weight measurement, mean of the volume measurement, and the number of the measurements, respectively.

To simply the analysis, it is assumed that the weight measurements are quite accurate, i.e., *wi* ≈ *w*0, therefore,

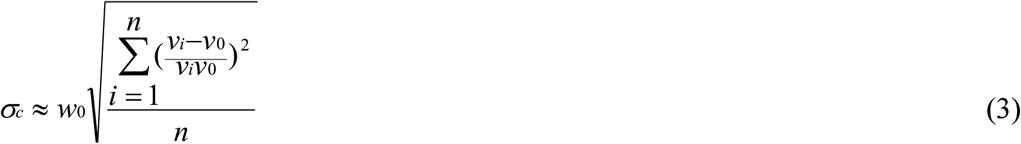

For a given shape of the volume,

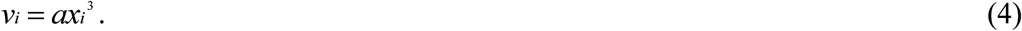

Therefore,

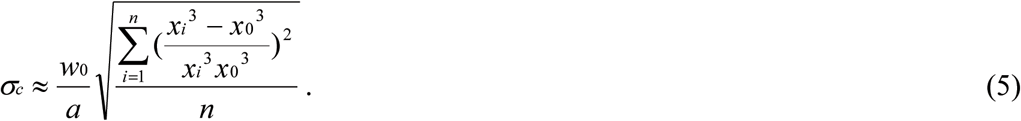

Because *xi*^3^ − *x*0^3^ ≈ *3xix0*(*xi* − *x*0) ≈ *3x0*^2^(*xi* − *x*0) when *xi* ≈ *x*0,

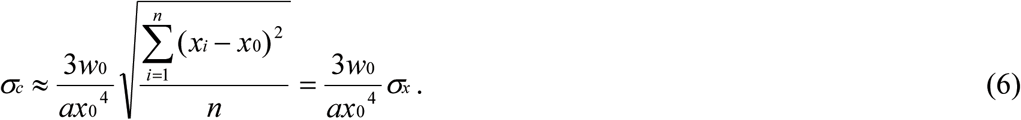

For a subcellular compartment, assuming its shape is a sphere, the above equation is then simplified as

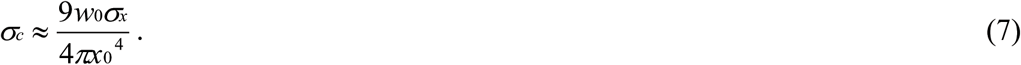

It is proposed that in Eq. 1, when *σx* = *σp*, it strikes a balance between the measurement accuracy of the position and that of the momentum of a particle. Under this condition,

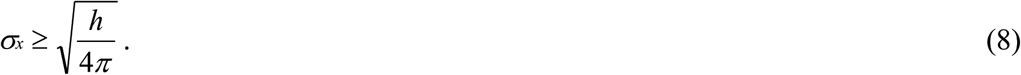

Assuming that *σx* in Eq. 7 is equivalent to *σx* in Eq. 8 since they both represent a standard deviation of distance measurements, and based on the early justification that the uncertainty level of a population of molecules is equal to or more than that of a subatomic particle,

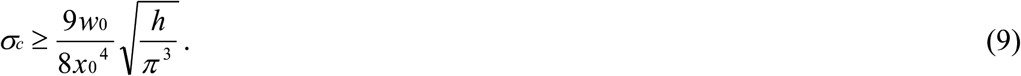

To calculate values of *σc* in Eq. 9, biologically relevant values of *w0* and *x0* should be determined. Based on the study by Milo and Phillips (2015), it is assumed that, in a subcellular compartment, *w0* = 4.9817 × 10^−18^g (the approximate weight of 100 molecules of a protein of 30000 Daltons), and *x*0 = 1 μm =10^−6^m, then

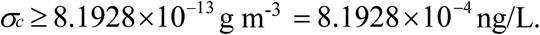

The above results indicate that when the *in vivo* concentration values are expressed in ng/L (likely being reasonably large absolute numerical values) in simulation of biochemical reactions, the relative numerical tolerance can be reasonably set at or around 10^−4^. For chemical concentrations expressed in a unit other than ng/L, the corresponding relative numerical tolerance values in simulation may be adjusted based on the above rvalue. The proposed range of relative numerical tolerance values suggests that the simulation results in Fig. 1A and B are likely closer to reality than that in Fig. 1C is.

Based on the above analysis, reducing relative numerical tolerance values for chemical concentration values in simulation will inevitably be met with increased standard deviation values in determining the energy (momentum) state of the molecules. This inverse relationship between the chemical concentrations and the momentum of the molecules likely means that the mathematical relationship among variables in a deterministic model will increasingly suffer inaccuracy with the increase in the accuracy of the concentration values. Furthermore, what is presented here may also be relevant in general to mathematical models that need to account for positions of molecules.

